# A Nonsecretory Antimicrobial Peptide Mediates Inflammatory Organ Damage in *Drosophila* Renal Tubules

**DOI:** 10.1101/2024.06.10.598165

**Authors:** Ayano Oi, Shun Nagashima, Natsuki Shinoda, Masayuki Miura, Fumiaki Obata

## Abstract

An excessive immune response damages organs, yet its molecular mechanism is incompletely understood. In this study, we used *Drosophila* renal tubules as a model to screen a factor mediating organ damage upon genetic activation of an innate immune Imd signalling pathway. We identified an antimicrobial peptide, Attacin-D (AttD), which causes organ damage upon Imd activation in the Malpighian tubules. Loss of AttD function suppresses most of the pathological phenotypes induced by Imd activation, such as cell death, compensatory stem cell proliferation, bloating of whole animal, susceptibility to a high salt diet, elevation of purine levels, and mortality, without compromising the immune activation. AttD is required for the immune-induced damage specifically in the Malpighian tubules but not the midgut. Interestingly, AttD uniquely lacks the signal peptide and is not secreted out from cells. Suppression of AttD almost completely attenuates mortality induced by gut tumour-induced immune activation. Our study elucidates the mechanistic effector of immune-induced organ damage.

## Introduction

Immune activation is a host defence mechanism that promotes tissue repair and eliminates pathogens such as bacteria or viruses, but excess immune activation harms host organs.^1^ It is widely accepted that chronic or prolonged inflammation is a common risk factor for many diseases, such as neurodegeneration, cardiovascular diseases, ischemic stroke, and cancer.^2–5^ However, as inflammation is a double-edged sword, general blockade of inflammatory pathways by immunosuppressive drugs often results in poor outcomes with side effects.^6,7^ To minimise the negative effect of the present clinical approach, it is necessary to know how immune activation leads to organ damage at the molecular level.

*Drosophila melanogaster* is a tractable model organism to identify the inflammatory organ damage mechanism due to the sophisticated genetic tools and evolutionally conserved innate immunity.^8^ *Drosophila* immunity is composed of several major signalling pathways, including immune deficiency (Imd) pathway. The Imd pathway resembles to the mammalian tumour necrosis factor receptor (TNFR) signalling pathway and is activated predominantly in response to bacterial peptidoglycan, which is detected by peptidoglycan recognition protein (PGRP) -LC or -LE.^9–11^ Signal transduction of Imd pathway leads to the proteolytic cleavage of the N-terminus of the Imd protein and subsequent activation of nuclear factor-kappa B (NF-κB) called Relish (Rel), which induces the expression of its target genes, such as antimicrobial peptide (AMP) genes.^9–11^ The Imd pathway is chronically activated during normal ageing and by necrotic cells genetically induced in wing cells, partly through gut microbiota.^12–14^ Suppression of Imd activation by overexpressing a negative regulator of the Imd pathway or removing gut microbiota can attenuate mortality and increase organismal lifespan.^12–15^ Chronic or excessive activation of the Imd pathway has been shown to induce tissue damage in various organs, such as the fat body, gut, and brain, in *Drosophila*.^16–18^ However, our knowledge of how an activated Imd pathway leads to tissue damage is limited, since Rel targets more than a hundred genes.^19^ If there is a specific factor that mediates tissue damage, loss of function of that gene should restore Imd-induced damage. To the best of our knowledge, such genes have not yet been identified.

Malpighian tubules (MTs) are the counterpart of mammalian renal tubules in *Drosophila*. There are two dominant cell types in the Malpighian tubules: larger principal cells and smaller stellate cells, with the former approximately four to five times more abundant than the latter^20^. Using these cells, MTs transport ions and water from or to the haemolymph to maintain fluid homeostasis by producing urine.^21^ Increased mortality from a high-sugar or a high-yeast diet is attributed to the decline in tubular function with stones in tubules composed of purine metabolites such as uric acid.^22,23^ Age-related Imd activation of MTs also leads to a shortened lifespan and an increase in purine levels.^24^ It has been reported that Imd activation in MTs induces a “bloating” phenotype, which indicates dysregulated water homeostasis during chronic oral infection,^25^ suggesting that infection-induced Imd activation causes a decline in renal tubular function. Inflammatory damage to MTs has also been found in a *Drosophila* cancer model. Overexpression of the active form of a transcription factor *yorkie* (*yki^3SA^)*, a homologue of the human oncogene, yes-associated protein 1 (YAP1), in intestinal stem cells is reported to induce tumour-derived pathology that includes activation of the Imd pathway in MTs, leading to a bloating phenotype and mortality.^26^ Homozygous mutation of *Rel* exacerbates the mortality of the cancer fly, but heterozygous *Rel* mutation or knockdown of *Rel* only in the MTs can partially restore it.^26^ The critical molecule, however, to induce Imd-dependent MT damage by the gut tumour is yet to be elucidated.

In this study, we first described how Imd activation in MTs leads to morphological, transcriptional, and organismal alterations. Then, we performed genetic screening to identify the Rel target genes in MTs responsible for Imd-induced organ damage. We identified Attacin-D (AttD) as a mechanistic target of Imd-induced pathological phenotypes, including bloating and mortality. Furthermore, loss of function of AttD restored mortality of the gut tumour model, suggesting the pathophysiological relevance of the gene for immune-induced tubular damage. Altogether, we report for the first time the essential effector of inflammatory organ damage in *Drosophila* renal tubules.

## Results

### Imd activation damages Malpighian tubules

To manipulate genes in MTs, we used two MT-specific Gal4 drivers, *Uro-Gal4* (expressed in principal cells of the main segment) and *NP1093-Gal4* (expressed in entire MTs) (Fig. 1A and S1A). We confirmed that *Uro-Gal4* was expressed only in principal cells, while *NP1093-Gal4* was expressed both principal and stellate cells (Fig. S1B). For *Uro-Gal4*, a temperature-sensitive form of Gal80 (*Uro-Gal4, tub-Gal80^ts^*, *Uro^ts^*) or GeneSwitch (*Uro^GS^*) is used to allow temporal gene manipulation in adult stages. GeneSwitch is a drug (RU486)-inducible system that enables us to negate the effect of genetic background. To activate the Imd pathway, we overexpressed the constitutive active form of Imd (*imd^CA^*), whose inhibitory N-terminus is removed.^27^

**Fig. 1.**
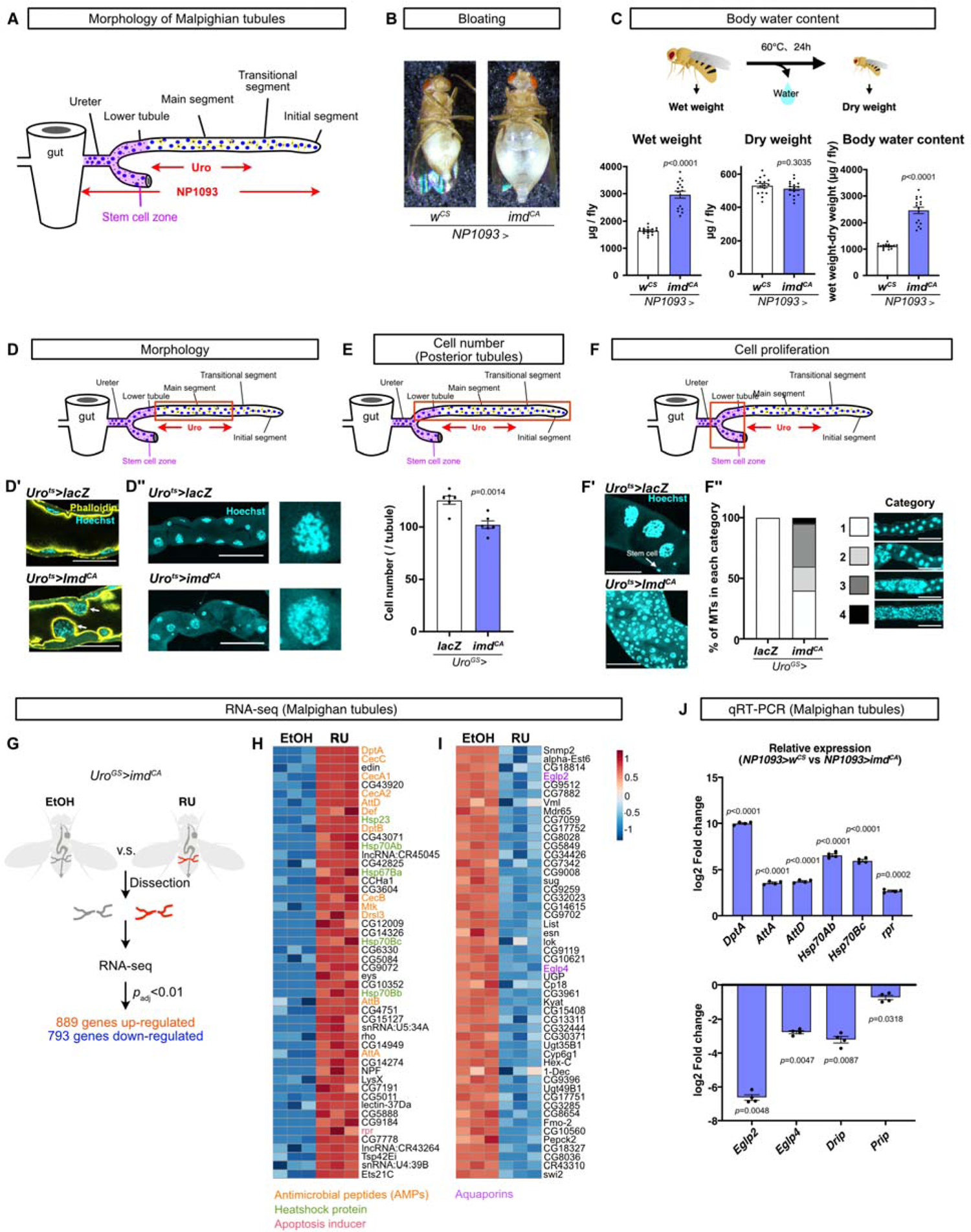
Imd activation damages Malpighian tubules. (A) A schematic image of the expression pattern of MT-specific drivers. (B, C) The bloating phenotype of 10-day-old female flies. A picture of each fly (B) and quantification of wet weight, dry weight, and body water content (C). n=16 for both genotypes. Statistics: unpaired two-tailed Student’s *t* test. (D-D’’) Representative images of the MTs (main segment) of female flies reared at 29 °C for 6 days. Cyan: Hoechst 33342 staining (nucleus), yellow: phalloidin staining (actin). Cross-sectional images, scale bar: 50 µm (D’) and Z-stacked images, scale bar: 100 µm (D’’). (E) Quantification of the cell number of posterior tubules in female flies fed a fly diet containing 50 µM RU486 for 12 days. (F-F’’) Representative images of stem cell proliferation. Z-stacked images of the MTs (stem cell zone) of female flies reared at 29 °C for 6 days, scale bar: 50 µm (F’). Quantification of the length of the area where stem cell proliferation was observed in female flies fed a fly diet containing 50 µM RU486 for 12 days (F’’). Stem cell proliferation was quantified by the length of the proliferated area: 1: length<0.1 mm, 2: 0.1 mm≤length<0.5 mm, 3: 0.5 mm≤length<1.5 mm, 4: 1.5 mm≤length. n=13 for *Uro^GS^>lacZ* and n=20 for *Uro^GS^>imd^CA^*. Scale bar: 500 µm. Statistics: unpaired two-tailed Student’s *t* test. (G) A schematic image of transcriptomic analysis. (H, I) Heatmap showing the top 50 genes with increased (H) or decreased (I) expression in the MTs of female flies fed a fly diet containing EtOH or 50 µM RU486 for 6 days. n=3 for both conditions. See Table S1 for detailed gene list and gene ontology analysis. (J) Quantitative RTLPCR of MTs of 10-day-old female flies, shown relative to control (*NP1093>w^CS^*). n=4 for both genotypes. Statistics: unpaired two-tailed Student’s *t* test. Each graph shows the mean ± SEM.

First, we overexpressed *imd^CA^* using *NP1093-Gal4* (*NP1093>imd^CA^*) to activate the Imd pathway in the whole organ. Adult female flies showed a typical “bloating” phenotype where the flies’ abdomen was swelled by excess water (Fig. 1B). This is consistent with a previous study showing that overexpression of wild type *imd* in MTs (using *c42-Gal4*), as well as chronic infection with *Ecc15*, caused bloating phenotype.^25^ We quantified body water content by measuring the body weight of each fly before and after drying (referred to as wet weight and dry weight, respectively) and found that Imd activation by *NP1093-Gal4* increased the wet weight compared to its isogenic background control *NP1093>w^CS^* (Fig. 1C). The Imd-activated flies did not show any difference in dry weight, suggesting that Imd activation increased water content without affecting development (Fig. 1C). When *imd^CA^* was overexpressed in the main segment of MTs using *Uro^ts^* or *Uro^GS^*, no bloating phenotype was observed. Under this condition, we conducted a Ramsay assay^28^ and confirmed the decline in the fluid secretion rate (Fig. S1C). These data indicate that Imd activation by overexpressing *imd^CA^* using both *NP1093* and *Uro* decreases tubular secretory function. The function of MTs is to maintain ion and water balance. To test the resistance to osmotic stress, we fed the flies high-NaCl food and measured the survival rate. As we expected, the survival rate of adult female flies on high-NaCl food was compromised by Imd activation by the two drivers (Fig. S1D). Previously, we showed that Imd activation by the *Uro* driver shortened lifespan and increased allantoin, a biomarker for purine levels in the body.^24^ We confirmed that Imd activation by *Uro^GS^*and *NP1093-Gal4* resulted in the same phenotypes (Fig. S1E and S1F). These data suggested that MT-specific Imd activation led to systemic pathological phenotypes.

Next, we observed the morphology of adult female MTs in which Imd was activated by *Uro^ts^* or *Uro^GS^* (*Uro^ts^>imd^CA^* or *Uro^GS^ >imd^CA^*). We occasionally noticed that some cells in the main segment fell into the lumen side (Fig. 1D and 1D’). The cells did not align well, and each nucleus tended to be larger and less condensed in the MTs with Imd activation than in the control (Fig. 1D’’). The number of cells of MTs was decreased (Fig. 1E), suggesting that principal cells underwent cell death. MTs are known to have stem cells in the ureter and lower tubules (Fig. 1A), which proliferate in response to tubular damage.^29^ We found that Imd activation in the main segment of MTs led to an increased number of small nuclei in the ureter and lower tubules (Fig. 1F, 1F’ and 1F’’). These data together suggested that Imd activation induced cell death and organ damage.

### Transcriptome analysis of Imd induced Malpighian tubules

To elucidate the molecular mechanism and to describe the transcriptional response of Imd-induced organ damage, we conducted RNA-seq analysis using dissected MTs with or without Imd activation (*Uro^GS^>imd^CA^*, EtOH (negative control) or RU486). We found 889 upregulated and 793 downregulated genes with adjusted *p* values less than 0.01 (Fig. 1G and Table S1). As expected, many immune-related genes were upregulated (Fig. 1H and Table S1). We noticed the upregulation of *heat shock proteins* (*Hsps*), suggesting that Imd activation induced a stress response in addition to classical NF-κB target genes (Fig. 1H).

Interestingly, two major aquaporins in principal cells, *Entomoglyceroporin 2* (*Eglp2*) and *Entomoglyceroporin 4* (*Eglp4*), were both downregulated (Fig. 1I). Although aquaporins in stellate cells, *Drip* and *Prip* did not change in this condition (Table S1), the deregulation of water homeostasis and consequent bloating phenotype might be partly due to the decline in the expression levels of aquaporins in principal cells. We also found that the expression of one of the apoptosis-inducible genes, *reaper* (*rpr*), was induced by Imd activation (Fig. 1H), indicating that the canonical apoptosis pathway was activated by Imd activation.

Upregulation of *AMPs*, *Hsps* and *rpr*, and downregulation of *Eglp2/4* were also observed in the MTs of *NP1093>imd^CA^* flies (Fig. 1J). Unlike *Uro^GS^>imd^CA^*, we found that *Drip* and *Prip* were downregulated (Fig. 1J), possibly due to the expression of *NP1093-Gal4* also in stellate cells. It has been shown that stellate cells transport water and chloride ions.^30–32^ The bloating phenotype observed only in *NP1093>imd^CA^* flies may be attributed to the decreased expression of these aquaporins in stellate cells as well as principal cells.

### Apoptosis is partly required for Imd-induced renal damage

Since *rpr* was induced by Imd activation, we hypothesised that apoptosis pathway was involved in the pathology. To investigate its contribution, first we directly activated apoptosis pathway. We used *Uro^GS^* to overexpress *rpr* in adult stage because overexpression of *rpr* using *NP1093-Gal4* resulted in developmental lethality. As expected, number of tubular cells decreased (Fig. 2A). Increased expression of *Hsps* suggested that the stress response is triggered by the tubular damage rather than direct target of Imd activation (Fig. 2B). We also found the vulnerability to NaCl diet (Fig. 2C) and increased systemic allantoin (Fig. 2D). These data suggested that induction of cell death of the principal cells is sufficient for induction of the pathological phenotypes seen in the flies with Imd activation in MTs.

**Fig. 2.**
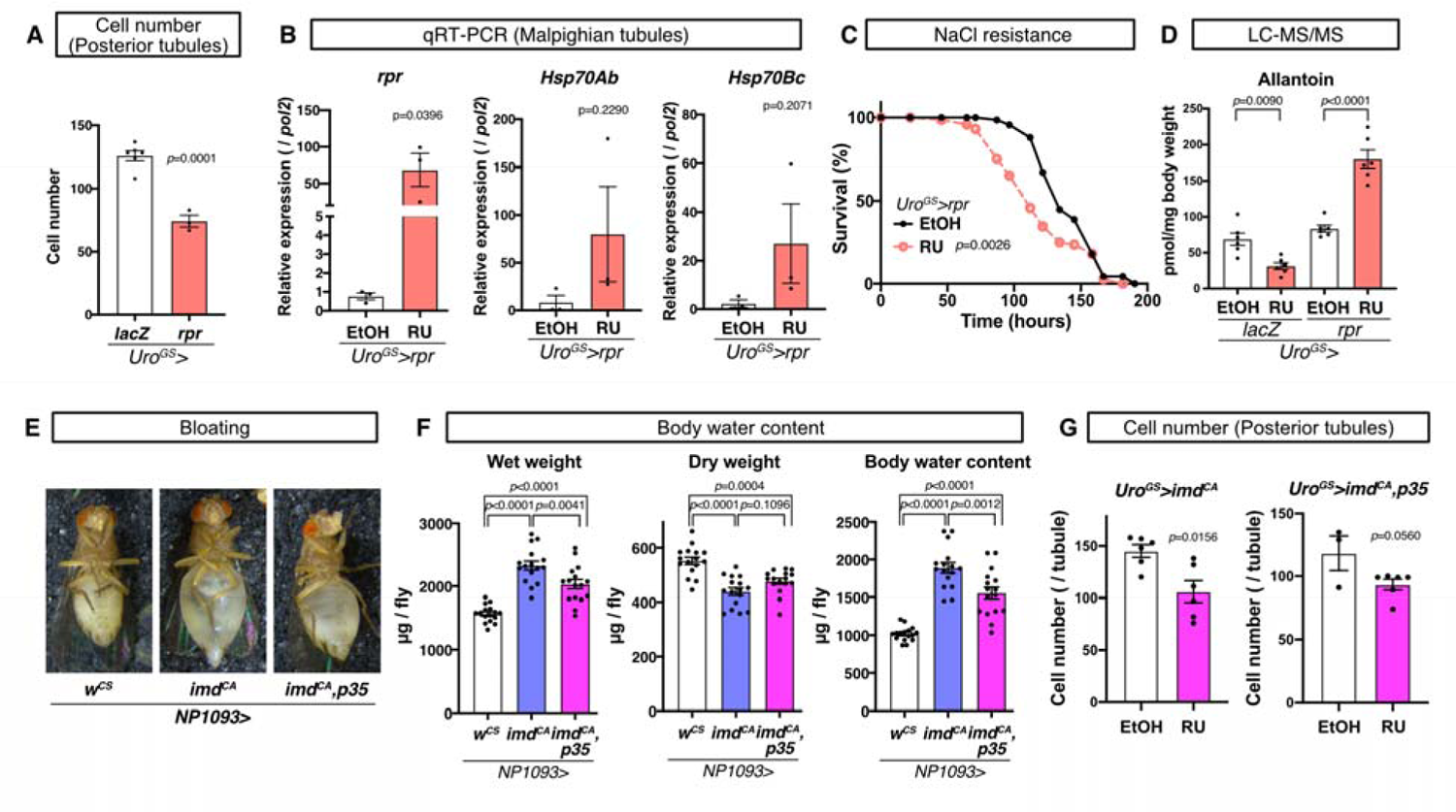
Apoptosis is partly required for Imd-induced renal damage. (A) Quantification of the cell number of posterior tubules of female flies. n=6 for *Uro^GS^>lacZ* and n=3 for *Uro^GS^>rpr*. Statistics: unpaired two-tailed Student’s *t* test. (B) Quantitative RTLPCR of MTs of female flies. n=3 for all genotypes. Statistics: unpaired two-tailed Student’s *t* test. (C) 500 mM NaCl resistance of female flies. n=67 for EtOH and n=72 for RU. Statistics: log-rank test. (D) Quantification of allantoin by LC-MS/MS in the whole body of female flies. n=6 for all genotypes. Statistics: one-way ANOVA with Sidak’s multiple comparisons. (E-F) The bloating phenotype of 10-day-old female flies. A picture of each fly (E) and quantification of wet weight, dry weight, and body water content (F). n=16 for all genotypes. Statistics: one-way ANOVA with Tukey’s multiple comparisons. (G) Quantification of the cell number of posterior tubules of female flies. n=3 for *Uro^GS^>imd^CA^, p35* EtOH and n=6 for all other genotypes. Statistics: unpaired two-tailed Student’s *t* test. For (A)-(D), and (G), female flies were fed a fly diet containing EtOH or 50 µM RU486 for 12 days before the experiments. Each graph shows the mean ± SEM.

Next, we examined if blocking apoptosis can restore the pathology of Imd activation by overexpressing a caspase inhibitor *p35*. We observed that the Imd-driven bloating phenotype and decreased cell number was partially suppressed (Fig. 2E-2G). These data indicate that apoptosis contributes to the pathology of Imd activation only partially and there should be additional mechanism of the inflammatory renal damage.

### *Attacin-D* is required for Imd-induced renal damage

Imd pathway has two main transcriptional regulators downstream, *Rel* and *AP1* (through the JNK pathway) (Fig. S2A). ^9^ To reveal which pathway contributed to organ damage, we genetically suppressed each pathway in the MTs with Imd activation. Suppressing the JNK pathway by overexpressing the inhibitor of JNK, *puckered* (*puc*), could not restore the decreased cell number and NaCl resistance (Fig. S2B and S2C), while knocking down *Rel* completely rescued the two phenotypes (Fig. S2B and S2C). These data led us to consider that *Rel* induces a hypothetical gene, “*<u>kil</u>ler of cells upon immune activation* (*Kili*)”, in the MTs that is responsible for renal damage observed in the flies with Imd activation.

To identify the *Kili* gene, we conducted RNAi screening using a list of genes upregulated by Imd-activated MTs (Fig. 3A and Table S1). We selected the top one hundred highly inducible genes as *Kili* candidates. We excluded noncoding RNAs or genes for which RNAi lines are unavailable in *Drosophila* stock centers, narrowing the number to 80 genes (Table S2). We crossed these knockdown lines with flies carrying both *Uro^GS^* and *Imd^CA^*, and manually observed the morphology of MTs, mainly focusing on stem cell proliferation as a proxy of tubular damage. Strikingly, we found that only one gene was necessary for Imd-induced renal damage, which was *Attacin-D* (*AttD*) (Fig. 3A), one of the β-barrel structure AMPs (Fig. S3A and Flybase) found in *Drosophila melanogaster* by genome sequence analysis.^33^ In *Drosophila melanogaster*, AMPs are divided into eight different families.^34^ *AttD* is one of the member of Attacin family (AttA, B, C, and D), but AttD has some unique features: 1) while other Attacin family genes are in the 2^nd^ chromosome, *AttD* is in the 3^rd^ chromosome, 2) the amino acid sequence of *AttD* is only 33% identical to *AttA,* while *AttB* is 98% and *AttC* is 73% identical to *AttA*, and 3) *AttD* lacks a signal peptide.^33,35^ Indeed, out of 42 AMP/AMP-like genes previously reported,^35–42^ we noticed the absence of a signal peptide sequence only in *AttD* predicted by SignalP 6.0 (Fig. S3B and Table S3).^43^ To observe the localisation of AttD *in vivo*, we created *AttD-sfGFP* knock-in allele by CRISPR/Cas9, in which Superfolder GFP (sfGFP) is fused to the C-terminus of the endogenous *AttD* (Fig. S3C). We confirmed that the *AttD-sfGFP* reporter fluorescence was induced by Imd activation (Fig. S3D), implying that it recapitulated the endogenous expression pattern as we expected. The signal of *AttD-sfGFP* was restricted to *Uro-Gal4* positive main segment, suggesting that AttD indeed remained in the cells and not secreted out of cells (Fig. S3D). Of note, the AttD-sfGFP knock-in flies should work as a good tool for monitoring Imd activation, as it reflects endogenous level of AttD which remains in the Imd-activated cells unlike other AMPs.

**Fig. 3.**
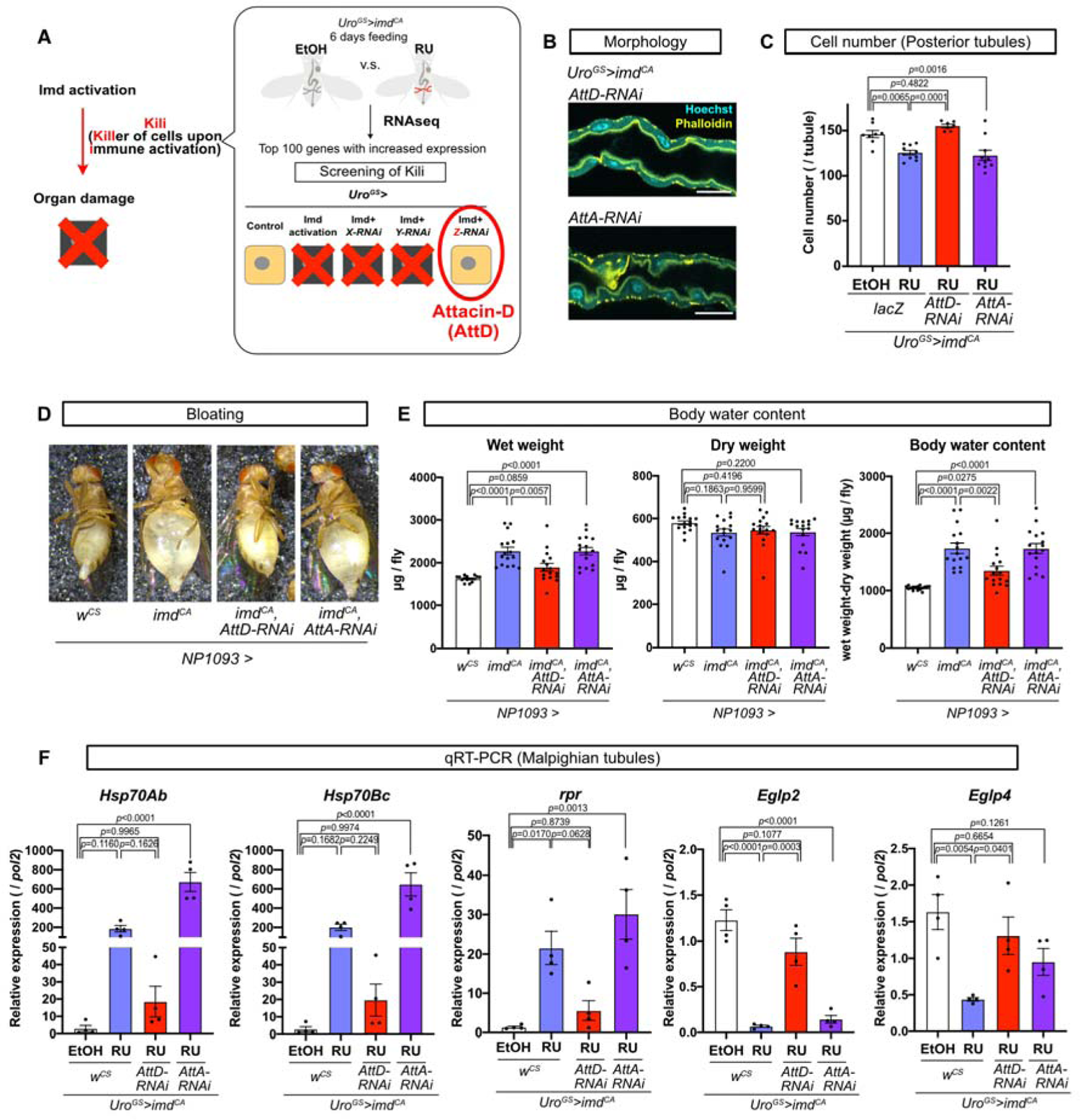
Genetic screening identifies Attacin-D as a crucial factor for Imd-induced renal damage. (A) A schematic image of genetic screening. (B) Representative images of the MTs (main segment) of female flies. Cyan: Hoechst 33342 staining (nucleus), yellow: phalloidin staining (actin), scale bar: 50 µm. (C) Quantification of the cell number of posterior MTs in female flies. n=8 for *Uro^GS^>imd^CA^, lacZ* EtOH, n=10 for *Uro^GS^>imd^CA^, lacZ* RU, n=7 for *Uro^GS^>imd^CA^, AttD-RNAi* RU, n=10 for *Uro^GS^>imd^CA^, AttA-RNAi* RU. Statistics: one-way ANOVA with Tukey’s multiple comparisons. (D-E) The bloating phenotype of 10-day-old female flies. A picture of female flies (D) and quantification of wet weight, dry weight, and body water content (E). n=16 for all genotypes. Statistics: one-way ANOVA with Tukey’s multiple comparisons. (F) Quantitative RTLPCR analysis of the MTs of female flies. n=4 for all genotypes. Statistics: one-way ANOVA with Tukey’s multiple comparisons. For (B), (C), and (F), female flies were fed a fly diet containing EtOH or 50 µM RU486 for 12 days. Each graph shows the mean ± SEM.

When we knocked down *AttD*, many phenotypes induced by Imd activation were rescued. Upon *AttD* knockdown, the disrupted morphology, and the decreased cell number in Imd-activated MTs were recovered (Fig. 3B and 3C). In sharp contrast, both phenotypes were not affected by knocking down *AttA* (Fig. 3B and 3C), which is one of the Attacin family members also upregulated by *imd^CA^*overexpression, suggesting the specificity of *AttD* in renal damage. We also observed that the bloating phenotype was totally suppressed by knocking down *AttD* but not *AttA* (Fig. 3D and 3E). The upregulation of *Hsp* and *rpr*, and the downregulation of *Eglp2/4* in the Imd-activated MTs were also rescued by *AttD* knockdown (Fig. 3F). Importantly, AMPs other than *AttD*, such as *DptA* and *AttA*, were still induced in this condition (Fig. S4A), indicating that the activity of Rel *per se* was not affected by *AttD-RNAi*. We confirmed that the decreased secretion rate of the dissected MTs was attenuated by *AttD-RNAi* (Fig. S4B). In addition, the increase in allantoin in the whole body and shortened lifespan of Imd-activated flies were restored (Fig. S4C and S4D), suggesting that systemic pathological phenotype could also be mediated by *AttD*.

### Mutation of *Attacin-D* supresses the Imd induced renal damage

To confirm the function of *AttD* in Imd-induced organ damage, we used a deletion mutant of *AttD*, named *AttD^SK1^*.^44^ In the *AttD^SK1^* mutant, 4 bases in the 546 bp *AttD* coding sequence are deleted, and a frameshift occurs to create a premature stop codon to produce a peptide with only 78 amino acids, leading to loss of function of *AttD*.^44^ As expected, loss of AttD function in the Imd-activated flies did not affect the upregulation of other AMPs, such as *DptA* and *AttA* (Fig. 4A). Activation of the Imd pathway in the *AttD^SK1/SK1^* failed to decrease the cell number in the organ (Fig. 4B). Accordingly, altered gene expression of *Hsps*, *rpr*, and *Eglp2/4* was restored in the *AttD* mutant (Fig. 4C). Consistently, elevation of whole-body allantoin levels by Imd activation was suppressed in the *AttD^SK1^* homozygous mutant (Fig. 4D). Strikingly, the shortened lifespan of flies with Imd activation in the MTs was almost completely restored by *AttD* mutant (Fig. 4E). Together, we identified AttD as a critical factor acting to induce tubular damage downstream of Imd activation.

**Fig. 4.**
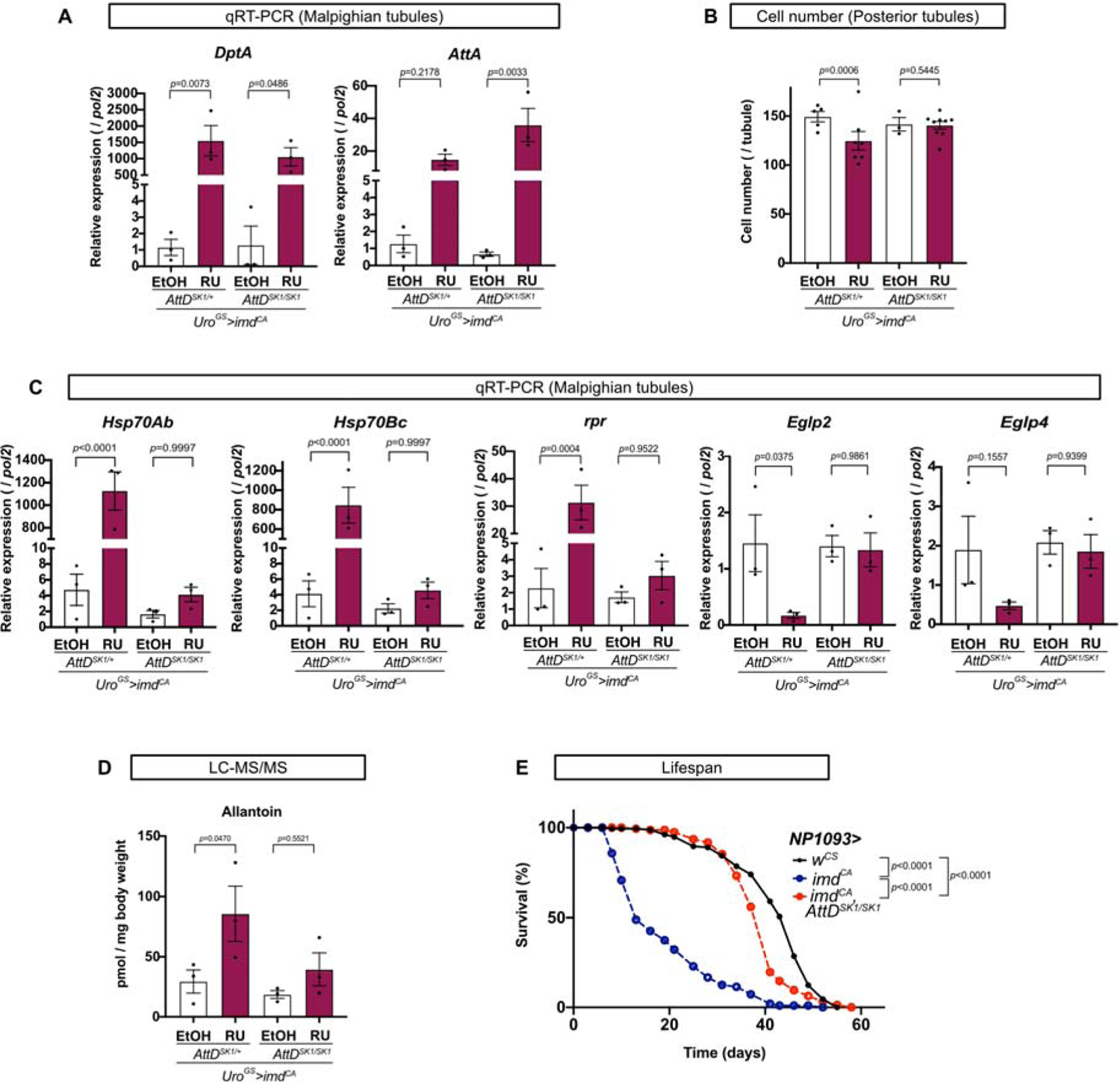
Mutation of *Attacin-D* suppresses Imd-induced renal damage. (A) Quantitative RTLPCR of MTs of female flies. n=3 for all genotypes. Statistics: one-way ANOVA with Sidak’s multiple comparisons. (B) Quantification of the cell number of posterior tubules of female flies. n=5 for *Uro^GS^>imd^CA^, AttD^SK1/+^*EtOH, n=7 for *Uro^GS^>imd^CA^, AttD^SK1/+^*RU, n=3 for *Uro^GS^>imd^CA^, AttD^SK1/SK1^*EtOH, and n=9 for *Uro^GS^>imd^CA^, AttD^SK1/SK1^*RU. Statistics: one-way ANOVA with Sidak’s multiple comparisons. (C) Quantitative RTLPCR of the MTs of female flies. n=3 for all genotypes. Statistics: one-way ANOVA with Sidak’s multiple comparisons. (D) Quantification of allantoin by LC-MS/MS in the whole body of female flies. n=3 for all genotypes. Statistics: one-way ANOVA with Sidak’s multiple comparisons. (E) Lifespan of female flies. n=169 for *NP1093>w^CS^*, n=113 for *NP1093>imd^CA^*, n=172 for *NP1093>imd^CA^, AttD^SK1/SK1^*. Statistics: log-rank test. For (A)-(D), female flies were fed a fly diet containing EtOH or 50 µM RU486 for 12 days before the experiments. Each graph shows the mean ± SEM.

### *Attacin-D* is not required for Imd-induced damage in other tissues

The discovery of AttD for immune-induced renal damage implied the possibility that the gene could mediate inflammatory damage in other parts of the body. To determine whether the function of AttD is generalised for other organs or specific to MTs, we tested the contribution of AttD in the adult gut and larval wing discs. In the adult midgut, Imd activation by infection has been shown to induce apoptosis.^16^ Consistent with this finding, qRTLPCR of the Imd-activated gut (*NP1^ts^>imd^CA^*) showed the induction of *rpr* and the damage-inducible ligand *Unpaired 3* (*Upd3*), which is known to stimulate stem cell proliferation in the gut (Fig. 5A). When we knocked down *AttD*, we still observed the upregulation of *rpr* and *Upd3* in the Imd-activated gut (Fig. 5A). In addition, we found that the lifespan shortening effect by *Imd^CA^*cannot be mitigated by *AttD-RNAi* in the gut (Fig. 5B), showing a clear contrast to what we observed in MTs (Fig. 4E and S4D).

**Fig. 5.**
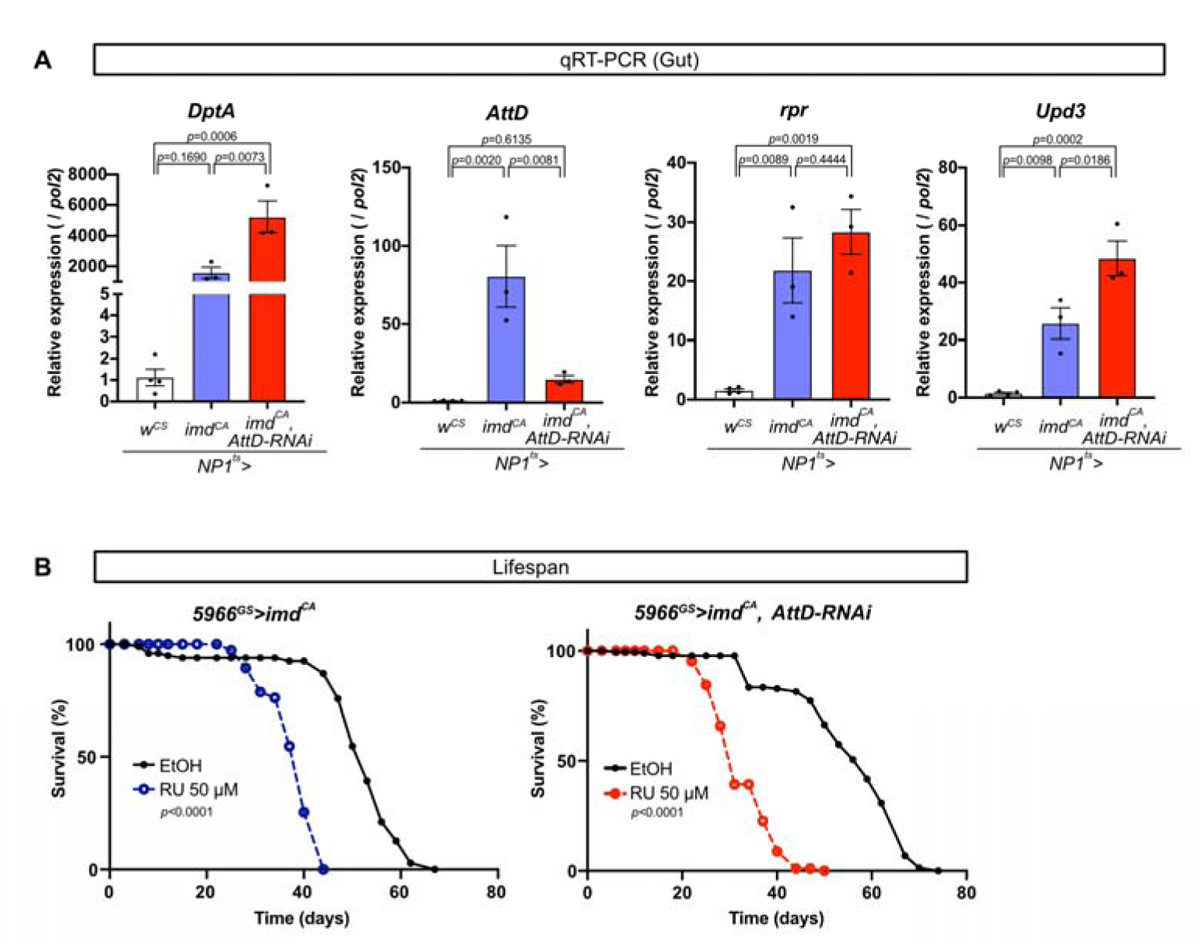
Attacin-D is not required for Imd-induced damage in the gut. (A) Quantitative RTLPCR of the gut of female flies reared at 29℃ for 7days. n=4 for *NP1^ts^>mCD8-PARP-VENUS, w^CS^*, n=3 for *NP1^ts^>mCD8-PARP-VENUS, imd^CA^*, and n=3 for *NP1^ts^>mCD8-PARP-VENUS, imd^CA^, AttD-RNAi*. Statistics: one-way ANOVA with Tukey’s multiple comparisons. (B) Lifespan of female flies fed a fly diet containing EtOH or 50 µM RU486. n=97 for *5966^GS^>imd^CA^*EtOH, n=97 for *5966^GS^>imd^CA^* RU, n=140 for *5966^GS^>imd^CA^, AttD-RNAi* EtOH, n=126 for *5966^GS^>imd^CA^, AttD-RNAi* RU, Statistics: log-rank test. Each graph shows the mean ± SEM.

We also tested the ability of AttD to induce cell death in the wing discs in larvae since they are frequently used as a model to study cell death machinery in *Drosophila*.^45^ *Rpr* overexpression by *WP-Gal4* strongly induced apoptosis in the wing pouch, which can be visualised by a cleaved PARP (cPARP) signal (Fig. S5A).^46^ Overexpressing *imd^CA^* also increased cPARP signal in the wing discs, although the phenotype was much milder than that of *rpr* (Fig. S5A and S5B). However, the increased cPARP signal by Imd activation did not decrease by knocking down *AttD* (Fig. S5C and S5D). From these results, we concluded that the function of AttD in causing damage by Imd activation could be specific to MTs.

### *Attacin-D* mediates tubular damage by gut tumour-induced Imd activation

Finally, we explored the pathophysiological contexts in which Imd-induced damage to MTs could be mediated by *AttD*. Overexpressing *yki^3SA^*in gut progenitor cells (*esg^ts^>yki^3SA^*) increases Imd activity in MTs and induces a bloating phenotype.^26^ It has been shown that the tumour also decreases the ovary size, a sign of cachexia induced by tumour-derived ligands *Upd3*, *PDGF- and VEGF-related Factor 1 (Pvf1)* and *Ecdysone-inducible gene L2 (ImpL2)*. Attenuation of the Imd pathway in the MTs does not suppress cachexia but partly rescues the overall mortality of the tumour fly.^26^ We therefore investigated whether the pathophysiology of this gut tumour model was *AttD* dependent.

As we expected, introduction of the *AttD^SK1^* mutation completely suppressed the bloating phenotype and the increased water content of *esg^ts^>yki^3SA^* without influencing tumour growth (Fig. 6A and 6B). We found that neither the decrease in ovary size nor the expression of tumour-derived cachectic ligands was restored in the *AttD* mutant (Fig. 6C and 6D), suggesting that cachexia is expectedly independent of the function of MTs. We also found that the gut tumour decreased flies’ NaCl resistance (Fig. 6E), suggesting that the flies are susceptible to osmotic stress. Furthermore, we found a decrease in *Eglp2/4* in an *AttD*-dependent manner (Fig. 6F). Strikingly, the survival of the gut tumour-induced flies was greatly improved in the *AttD* mutant in both females and males (Fig. 6G). These data suggest that AttD can be a crucial factor in determining the mortality of flies bearing gut tumours.

**Fig. 6.**
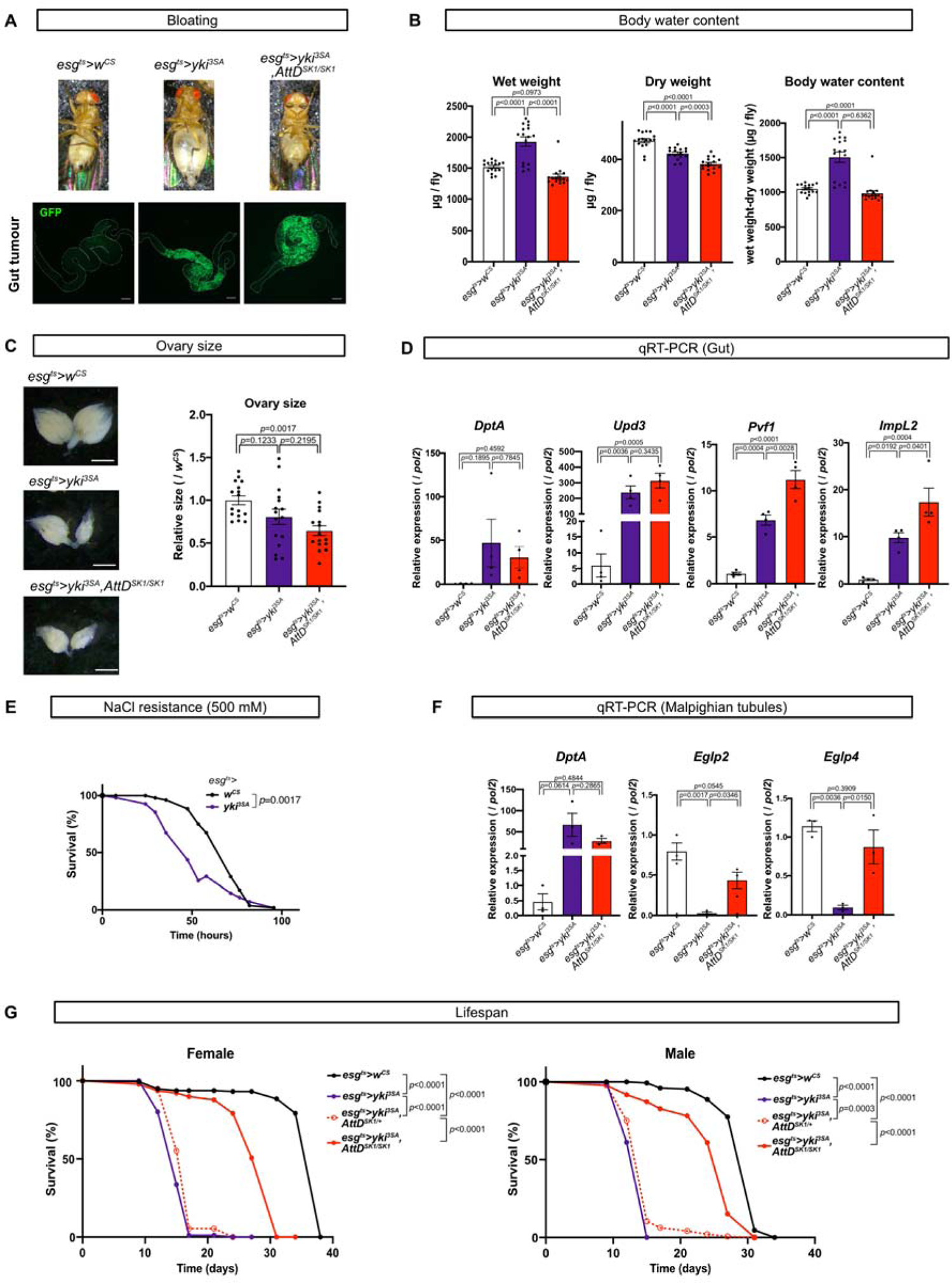
Attacin-D mediates renal damage and mortality in the gut tumour-bearing flies. (A) The bloating phenotype of female flies. Pictures of each fly (upper panel) and gut tumour (lower panel, GFP). Scale bar: 200 µm. (B) Quantification of wet weight, dry weight, and body water content. n=16 for all genotypes. Statistics: one-way ANOVA with Tukey’s multiple comparisons. (C) Representative images of ovaries and quantification of the ovary size of each genotype. Scale bar: 500 µm. n=16 for all genotypes. Statistics: one-way ANOVA with Tukey’s multiple comparisons. (D) Quantitative RTLPCR of the guts of female flies. n=4 for all genotypes. Statistics: one-way ANOVA with Tukey’s multiple comparisons. For (A)-(D), genes were manipulated at 29 °C for 8 days. (E) 500 mM NaCl resistance of female flies incubated at 29 °C. n=51 for *esg^ts^>w^CS^*and n=54 for *esg^ts^>yki^3SA^*. Statistics: log-rank test. (F) Quantitative RTLPCR of MTs of female flies reared at 29 °C for 10 days. n=3 for all genotypes. Statistics: one-way ANOVA with Tukey’s multiple comparisons. (G) Lifespan of female and male flies reared at 29 °C. n=175 for female *esg^ts^>w^CS^*, n=176 for female *esg^ts^>yki^3SA^*, n=176 for female *esg^ts^>yki^3SA^, AttD^SK1/+^*, n=104 for female *esg^ts^>yki^3SA^, AttD^SK1/SK1^*, n=144 for male *esg^ts^>w^CS^*, n=172 for male *esg^ts^>yki^3SA^*, n=144 for male *esg^ts^>yki^3SA^, AttD^SK1/+^*, and n=133 for male *esg^ts^>yki^3SA^, AttD^SK1/SK1^*. Statistics: log-rank test. Each graph shows the mean ± SEM.

## Discussion

The mechanism of how immune activation damages organs and hence induces organismal mortality has been intensively studied. Given that there are hundreds of immune-inducible genes, discovering the direct effector that causes inflammatory organ damage is challenging. From the genetic screening, our present study identified a single gene, *AttD*, acting as a host-damaging factor induced by the Imd pathway in *Drosophila* renal tubules. When *AttD* is mutated in Imd-activated MTs, functional and morphological defects are no longer observed, although other immune-inducible genes, such as AMPs, are still induced. Furthermore, loss of function of *AttD* can almost completely suppress the bloating phenotype and shortened lifespan of a fly with the gut tumour. To the best of our knowledge, *AttD* is the first gene that is involved in Imd activation-driven renal damage. The possibility that AMPs are responsible for immune-induced damage in other organs has been implicated in prior works. For example, overexpression of AMPs can shorten lifespan and induce cell death in neurons or fat bodies.^47,48^ However, the requirement of each AMP in immune-induced organ damage has not been fully addressed since AMPs are believed to work redundantly and synergistically.^44^ Recently, knocking down a single AMP, Metchnikowin (Mtk), has been shown to partially rescue poly (GR)-induced neurotoxicity, although how Mtk exacerbates cell damage has not been identified.^49^

The finding that apoptosis was only partially related to imd induced cell death led us to consider the existence of other type of cell death. Generally, AMPs are known to kill bacteria to permeabilise their membrane.^50^ Also, AttD has the β-barrel structure (Fig. S3A), which is known to form pores frequently in the mitochondrial membrane.^51^ Given that AttD cannot be secreted from cells, it is very likely that AttD can form pores to organelle membranes, including mitochondria, which is structurally similar to an ancient bacterium within cells. During pyroptosis, pore formation in plasma membrane is executed by oligomerisation of the N-terminal cleaved fragments of Gasdermin D.^52^ The plasma membrane is subsequently ruptured by cell-surface protein Ninjurin-1through oligomerisation.^53–55^ It is possible that AttD oligomerises to form pores. The structural investigation on how AttD kills cell can be the next fascinating question. Besides, the further elucidation of the mechanisms how and why AttD damages MTs, but not other organs, is necessary for the complete understanding.

Since our study only reveals the detrimental effects of AttD in pathological conditions during excessive and chronic immune activation, AttD’s physiological function remains unclear. Considering that AttD is the only AMP that cannot be secreted, AttD might be essential to kill intracellular pathogens. In *Drosophila*, *Listeria monocytogenes* is one of the bacteria that invades cells and induces Imd activation.^56^ One AMP called Listericin has already been found to combat *L. monocytogenes*, although it has a signal peptide.^41^ In mammals, *Legionella pneumophila* is a well-known intracellular bacterial pathogen that hijacks the early endosome and forms vacuoles to proliferate in host cells.^57^ The ER vesicular soluble N-ethylmaleimide-sensitive factor attachment protein receptors (v-SNAREs) are recruited to the vesicle.^58,59^ Interestingly, AttD is predicted to interact with SNARE proteins (^60^ and Flybase), suggesting that AttD might be important to combat such bacteria. Because *L. pneumophila* is reported to infect *Drosophila* cells,^61^ it would be interesting to study whether AttD shows the same localisation as *L. pneumophila* and whether AttD can kill them.

Tackling inflammatory diseases by suppressing immune signalling *per se* is a straightforward but harsh strategy, as immune signalling has both beneficial and detrimental aspects. Not surprisingly, homozygous null mutation of *Rel* in inflammatory fly models, such as flies with gut tumours, exacerbates mortality.^26^ In clear contrast, the *AttD* null mutation almost fully suppresses the mortality of the same cancer model. Therefore, the identification of a single effector, such as AttD, is crucial for clinical application. As AttD is specifically required for renal damage, other genes responsible for each cell type in other organs should be discovered. To date, no mammalian orthologues of *AttD* have been found, but we assume that structural or functional orthologues are present in mammals that may target the organelle membranes. It should become a prominent target for pharmacological intervention against human inflammatory diseases, including cancer.

## Acknowledgments

We thank Dr. Corey Goodman, Dr. Edan Foley, Dr. Norbert Perrimon, Dr. Wei Song, Dr. Julien Royet, the Kyoto Stock Center, National Institute of Genetics, Vienna Drosophila Resource Center and Bloomington Drosophila Stock for Drosophila stocks; Dr. Shigenobu Yonemura, Mr. Kenta Onoue, and Ms. Satoko Okayama for technical support; all members of Miura’s lab and Obata’s lab for technical assistance and important advice. This work was supported by AMED-PRIME to F.O. (JP20gm6310011), The Japan Society for the Promotion of Science to F.O. (22H02769) and to A.O. (23KJ1182), The Naito Foundation to F.O, and RIKEN Junior Research Associate Program to A.O.

## Author contributions

F.O. and A.O. conceived the project; A.O. performed most of the experiments and analysed the data; S.N. and N.S. helped bioimaging and data analysis; A.O., S.N., N.S., and F.O. wrote the initial manuscript; M.M. and F.O. supervised the study; All authors edited and approved the final manuscript.

## Methods

### *Drosophila* stock and husbandry

Flies were reared on a standard diet containing 4.5% cornmeal (NIPPN CORPORATION), 4% dry brewer’s yeast (Asahi Breweries, HB-P02), 6% glucose (Nihon Shokuhin Kako), and 0.8% agar (Ina Shokuhin Kogyo, S-6) with 0.4% propionic acid (Wako, 163-04726) and 0.03% butyl p-hydroxybenzoate (Wako, 028-03685). Flies were reared at 25 °C and 65% humidity with 12 h/12 h light/dark cycles.

The fly lines used in this study were as follows: *w^CS^*(*w^iso^*^31^ backcrossed with *Canton-S* eight times), *w^iso31^*, *NP1093-Gal4* (Kyoto Drosophila Stock Center, 103880), *Uro-Gal4, tub-Gal80^ts^*,^62^ *Uro-GeneSwitch*,^63^ *NP1-Gal4,tub-Gal80^ts^* (gifted from Dr. N. Perrimon), *5966-GeneSwitch* (gifted from Dr. H. Jasper)^64^, *esg-Gal4, tub-Gal80^ts^*,^64^ *UAS-lacZ* (gift from Dr. C. S. Goodman), *UAS-imd^CA^*,^27^ *UAS-2xeGFP* (Bloomington Drosophila Stock Center, 6874), *UAS-GFP.nls* (Bloomington Drosophila Stock Center, 4775), *UAS-p35* (Kyoto Drosophila Stock Center, 108018), *UAS-rpr* (Bloomington Drosophila Stock Center, 5824), *UAS-puc*^65^, *UAS-Relish-RNAi* (Bloomington Drosophila Stock Center, 33661), *UAS-AttD-RNAi* (Bloomington Drosophila Stock Center, 55979), *UAS-AttA-RNAi* (Bloomington Drosophila Stock Center, 56904), *AttD^SK1/SK1^*,^44^ *UAS-mCD8-PARP-VENUS*,^66^ *UAS-TdTomato-Sec61*β (Bloomington Drosophila Stock Center, 64746), *UAS-Bip-sfGFP-HDEL* (Bloomington Drosophila Stock Center, 64749), *UAS-mitoRFP* (Kyoto Drosophila Stock Center, 117016), *UAS-yki^3SA^* (Bloomington Drosophila Stock Center, 28817), *NP1093-Gal4*, *Uro-GeneSwitch*, *UAS-lacZ*, *UAS-imd^CA^*, and *AttD^SK1/SK1^* have backcrossed eight generations with *w^Dahomey^* or *w^CS^*. Fly lines used for genetic screening (Fig. 2A) are listed in Table S2.

Embryos were collected on an acetic acid agar plate (3% agar (Wako, 010-15815), 10% sucrose (Wako, 196-00015), and 0.35% acetic acid (Wako, 017-0256)) with live yeast paste (Oriental, 01402017) using a cage containing virgin female and male flies. Equal volumes of embryos were placed on the fly food. Eclosed flies were maintained for 2 days for maturation, and they were sorted by sex and put into vials (30 flies per vial). For manipulation using GeneSwitch, 50 µM RU486 (Tokyo Chemical Industry, M1732) dissolved in EtOH (Wako, 057-00451) or an equal volume of EtOH (as a control) was mixed in standard food.

### Generation of transgenic flies

To generate *AttD-sfGFP* knock in flies, we used CRISPR/Cas9 system to insert Superfolder GFP (sfGFP) at C terminus of the *AttD* gene (Kina et al., 2019). We used pScarlessHD-sfGFP-DsRed (Addgene, #80811) to distinguish successful insertion by eye fluorescence. An sgRNA target site of AttD was selected using CRISPR Optimal Target Finder (Gratz et al., 2014). Complementary oligonucleotides with overhangs were annealed and cloned into the BbsI-digested U6b vector using a DNA ligation kit (Takara Bio, 6023). Sense strand:TTCGGAAGTGCCAATCATCACCTC, anti-sense strand: AAACGAGGTGATGATTGGCACTTC. The targeting vector was constructed by inserting sfGFP-3×P3-DsRed with 500bp homology arms with a mutation in the PAM sequence next to the sgRNA site (from TGG to TAG). First, sfGFP-3×P3-DsRed and homology arms were PCR-amplified using PrimeSTAR^®^ Max DNA Polymerase (Takara Bio, R045). Primers for PCR were designed using the NEBuilder Assembly Tool. The gel-purified PCR products were cloned into the EcoRI-digested pBluescript II SK (+) vector using NEBuilder HiFi DNA Assembly Master Mix (New England Biolabs, E2621X). The mixture of *pU6b-sgRNA* and targeting vector was microinjected into *w^11^*^18^*; attP40{nos-Cas9}/CyO* embryos by WellGenetics. The F0 adults were crossed with balancer lines, and the sfGFP-inserted lines were selected by 3×P3-DsRed. After selection, 3×P3-DsRed was removed by crossing with *hsp70-PBac* (Bloomington Drosophila Stock Center 32174) and F1 flies without eye fluorescence were selected. The primers used for PCR are listed in Table S4.

### Quantification of body water content

The body weight of each fly was measured using a microbalance (METTLER TOLEDO, XR2UV), and they were defined as wet weight. To trace them, each fly was placed in a 0.2 ml tube one by one and sealed with Parafilm with a small air hole. After incubation at 60 °C for 24 hours to dry up, their body weight was measured again, and they were defined as dry weight. Body water content was calculated by subtracting the dry weight from the wet weight.

### Imaging of Malpighian tubules and quantification of morphology

Malpighian tubules were dissected in PBS and fixed with 4% PFA (Nacalai Tesque, 09154-14) for 30-60 minutes. They were washed with PBST (PBS containing 0.1% Triton X-100 (Nacalai Tesque, 35501-15)) and incubated with 0.8 µM Hoechst 33342 (Invitrogen, H3570) and Phalloidin 645 (1:10000, Abcam, AB176759) for 30-60 minutes at room temperature.

They were washed with PBST and mounted in SlowFade Gold (Invitrogen, S36936). Confocal images were obtained using a Leica SP8. For the quantification of cell number, all Hoechst signals in tubules other than small dots of stem cells were counted manually. For the quantification of stem cell proliferation, the length of the proliferated area was measured and classified as follows: 1: length<0.1 mm, 2: 0.1 mm≤length<0.5 mm, 3: 0.5 mm≤length<1.5 mm, and 4: 1.5 mm≤length. ZEISS Axio Vert. A1 was used for imaging.

To observe the expression pattern of *Uro-Gal4* and *NP1093-Gal4*, dissected MTs were fixed with 4% PFA for 60 minutes. They were washed with PBST, blocked in PBST with 5% normal donkey serum (PBSTn), and incubated with mouse anti-dlg monoclonal antibody (1:100, Developmental Studies Hybridoma Bank, #4F3, RRID: AB_ 528203) in PBST overnight at 4 °C. The samples were then washed with PBST, incubated at room temperature for 2 hours with Alexa Fluor® 647-conjugated donkey anti-mouse IgG (H+L) antibodies (1:500, Invitrogen, # A-31571, RRID: AB_162542) and 0.8 µM Hoechst 33342 at room temperature. They were washed with PBST and mounted in SlowFade Gold. Images were captured using a Leica TCS SP8 microscope (Leica Microsystems, Wetzlar, Germany).

To observe the localisation of AttD using knock in *AttD-sfGFP*, dissected MTs were incubated with 100nM MitoTracker^TM^ Deep Red FM (Thermo Fisher Scientific, M22426) for 2 hours at room temperature. After briefly washing in PBS, they were mounted in SlowFade Gold. Images were obtained using a Zeiss LSM880 and processed by Airyscan.

### Secretion assay

The tubular secretion assay was conducted as previously described.^67^. Briefly, anterior Malpighian tubules were carefully cut at the ureter in 1:1 *Drosophila* saline: Schneider’s medium. One side of the tubules was suspended in a 10-20 µl droplet of *Drosophila* saline containing 0.2% brilliant blue FCF (Wako, 027-12842), while the other side was wrapped around a metal pin. They were covered with fluid paraffin (Wako, 164-00476) to prevent evaporation. A schematic diagram of the experimental setup is shown in Fig. S1B. After 30-60 minutes of preincubation, excreta from the ureter were removed using a glass capillary, and the assay was started. Every 10 minutes, excreta were removed after the size was measured. The volume of secreted droplets was calculated to give a secretion rate (nl/min).

### NaCl resistance

Thirty adult flies per vial were put into a vial containing 1% agar (Wako, 010-15815), 5% sucrose (Wako, 196-00015), and 500 mM NaCl (Wako, 191-01665). Dead flies were counted several times each day.

### Lifespan measurement

Thirty adult flies per vial were placed in a vial containing a fly diet. Dead flies were counted every 2 to 4 days when the flies were transferred to fresh vials.

### Measurement of allantoin levels

Four whole bodies of adult female flies were homogenised in 160 µl of 80% methanol (Wako, 132-06471) containing 10 µM internal standards (methionine sulfone (Sigma, M0876-1G) and 2-morpholinoethanesulfonic acid (Dojindo, 341-01622)). Samples were centrifuged at 15000 ×g for 5 minutes, and the supernatant was collected. To remove proteins, 75 µl acetonitrile (Wako, 015-08633) was added and centrifuged at 15000 ×g for 5 minutes. After passing through a Nanosep 10 kDa column (PALL, OD010C35) by centrifugation at 14000 ×g for 10 minutes, the samples were completely evaporated. They were dissolved in ultrapure water (Invitrogen, 10977-015) and injected into an LC-MS/MS (Shimadzu, LCMS-8050) with a PFPP column (Sigma, Discovery HS F5, 2.1 mm × 150 mm, 3 µm) in a column oven at 40 °C. A gradient from solvent A (0.1% formic acid in water) to solvent B (0.1% acetonitrile) was performed for 20 minutes. MRM methods for metabolite quantification were optimised using the software (Shimadzu, LabSolutions).

### RNA-sequencing analysis for transcriptomics

RNA was extracted from 15 adult female MTs using a Promega ReliaPrep^TM^ RNA Miniprep Kit (Z6112). A cDNA library was prepared using the QuantSeq 3’ mRNA-Seq Library Prep Kit for Illumina (FWD) (Lexogen, 015.384). Sequencing was performed using Illumina NextSeq 500 and NextSeq 500/550 High Output Kit v.2.5 (75 cycles) (Illumina, 20024906). Raw reads were analysed by the BlueBee Platform (Lexogen), which performs trimming, alignment to the *Drosophila* genome and counting of the reads. The count data were analysed statistically by Wald’s test using DESeq2. Gene ontology analysis was done by DAVID tools. ^68,69^ The results have been deposited in DDBJ under the accession number DRA016964.

### Quantitative RTDPCR analysis

RNA was extracted from 5-6 female MTs or 3 female guts using a Promega ReliaPrep^TM^ RNA Miniprep Kit (Z6112). Quantitative RTLPCR was performed using a OneTaq RTLPCR kit (Promega, M0482S) with qTOWER3 (Analytik Jena). Pol2 was used as an internal control. Quantification was performed by the ΔΔct method. Primer sequences are listed in Table S4.

### Prediction of signal peptides

Amino acid sequences of AMP and AMP-like genes were obtained from FlyBase (https://flybase.org/). These sequences were submitted to the SignalP 6.0 website (https://services.healthtech.dtu.dk/services/SignalP-6.0/) and graphs showing signal peptide prediction were obtained.

### Imaging of wing discs and quantification of c-PARP signal

Wing imaginal discs were dissected in PBS and fixed with 4% PFA for 20 minutes. They were washed with PBST, blocked with PBSTn, and incubated with rabbit anti-cleaved PARP (Asp214) (D64E10) XP monoclonal antibody (1:100, Cell Signalling Technology, #5625, RRID: AB_10699459) in PBSTn overnight at 4 °C. The samples were then washed with PBST, incubated at room temperature for 2 hours with Cy3-conjugated donkey anti-rabbit IgG (H+L) antibodies (1:500, Jackson ImmunoResearch Labs, # 711-165-152, RRID: AB_2307443) and 8 µM Hoechst 33342 suspended in PBSTn, and washed again with PBST. Samples were mounted in SlowFade Gold. Images were captured using a Leica TCS SP8 microscope. For quantification, using Fiji software,^70^ the VENUS-positive area was enclosed with polygon tools, and the mean intensity of the c-PARP signal in the area was calculated.

### Imaging of ovaries and quantification of ovary size

Ovaries were dissected in PBS and lined up on glass slides. Images were captured using Leica MZ10F. For quantification, using Fiji software,^70^ the outline of the ovary was enclosed with polygon tools, and the size of the enclosed area was calculated.

### Quantification and statistical analysis

Statistical analysis was performed using GraphPad Prism 8 except for survival curves where OASIS2 was used.^71^ A two-tailed Student’s *t* test was used to test between two samples.

One-way ANOVA with Tukey’s multiple comparisons was used to compare the mean of each column with the mean of every other column. One-way ANOVA with Sidak’s multiple comparisons was used to compare the mean of selected pairs of columns. Dunnett’s multiple comparison was used to compare the mean of each column with the mean of a control column. The log-rank test was used to compare two survival curves. All the statistical details can be found in the figure legends. Bar graphs were drawn as the mean and SEM with all the data points shown by dots to allow readers to see the number of samples and each raw data point.

**Table S1** Differentially expressed genes in the adult MTs upon Imd activation (adjusted *p*-value<0.01), related to Fig. 1.

**Table S2** RNAi line list for genetic screening, related to Fig. 3.

**Table S3** Signal peptide and cleavage site predicted by SignalP 6.0, related to Fig. S3B. Table S4 Primer list, related to Methods.

