## Supplementary Figures for "A Nonsecretory Antimicrobial Peptide Mediates Inflammatory Organ Damage in *Drosophila* Renal Tubules"

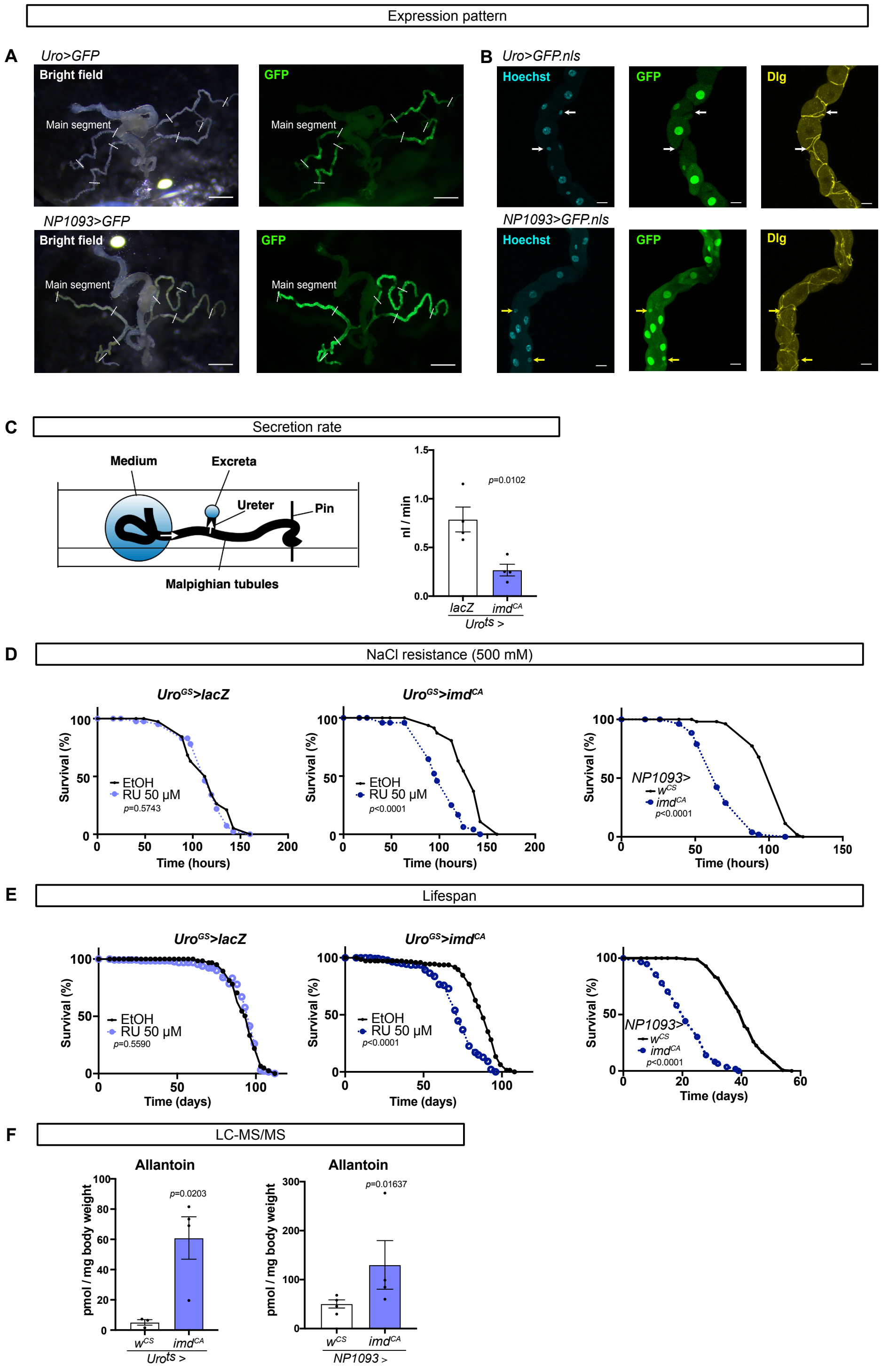

Expression pattern
A
Uro>GFP
Bright field
Main segment
GFP
Main segment
NP1093>GFP
Bright field
Main segment
GFP
Main segment
B
Uro>GFP.nls
Hoechst
GFP
Dlg
NP1093>GFP.nls
Hoechst
GFP
Dlg
C
Secretion rate
Medium
Excreta
Ureter
Pin
Malpighian tubules
nl / min
p=0.0102
lacZ
imdCA
Urots >
D
NaCl resistance (500 mM)
UroGS>lacZ
Survival (%)
EtOH
RU 50 μM
p=0.5743
Time (hours)
UroGS>imdCA
EtOH
RU 50 μM
p<0.0001
Time (hours)
NP1093>
wCS
imdCA
p<0.0001
Time (hours)
E
Lifespan
UroGS>lacZ
Survival (%)
EtOH
RU 50 μM
p=0.5590
Time (days)
UroGS>imdCA
EtOH
RU 50 μM
p<0.0001
Time (days)
NP1093>
wCS
imdCA
p<0.0001
Time (days)
F
LC-MS/MS
Allantoin
pmol / mg body weight
p=0.0203
wCS
imdCA
Urots >
Allantoin
pmol / mg body weight
p=0.01637
wCS
imdCA
NP1093 >

**Fig. S1 Imd activation in Malpighian tubules leads to systemic pathological phenotypes.**

(A) Expression pattern of the MT-specific divers *Uro-Gal4* (upper) and *NP1093-Gal4* (lower). Scale bar: 500 µm. (B) Representative Z-stacked images of the MTs (main segment) of female flies. Cyan: Hoechst 33342 staining (nucleus), Green: GFP, yellow: Dlg staining. Scale bar: 20 µm. White allows show stellate cells without GFP signal and yellow allows show stellate cells with GFP signal. (C) Quantification of the tubular secretion rate measured by Ramsay assay. The left panel is a schematic image of Ramsay assay. Female flies were reared at 29 °C for 7 days. n=4 for both genotypes. Statistics: unpaired two-tailed Student's *t* test. (D) 500 mM NaCl resistance of female flies. n=38 for $Uro^{GS}$>*lacZ* EtOH, n=41 for $Uro^{GS}$>*lacZ* RU, n=46 for $Uro^{GS}$>$imd^{CA}$ EtOH, n=48 for $Uro^{GS}$>$imd^{CA}$ RU, n=53 for *NP1093*>$w^{CS}$, and n=51 for *NP1093*>$imd^{CA}$. $Uro^{GS}$>*lacZ* and $Uro^{GS}$>$imd^{CA}$ flies were fed a fly diet containing EtOH or 50 µM RU for 12 days before the experiment. Statistics: log-rank test. (E) Lifespan of female flies. n=138 for $Uro^{GS}$>*lacZ* EtOH, n=130 for $Uro^{GS}$>*lacZ* RU, n=145 for $Uro^{GS}$>$imd^{CA}$ EtOH, n=141 for $Uro^{GS}$>$imd^{CA}$ RU, n=170 for *NP1093*>$w^{CS}$, and n=173 for *NP1093*>$imd^{CA}$. $Uro^{GS}$>*lacZ* and $Uro^{GS}$>$imd^{CA}$ flies were fed a fly diet containing EtOH or 50 µM RU. Statistics: log-rank test. (F) Quantification of allantoin by LC-MS/MS in the whole body of female flies. $Uro^{ts}$>*lacZ* and $Uro^{ts}$>$imd^{CA}$ flies were reared at 29 °C for 6 days, and *NP1093*>$w^{CS}$ and *NP1093*>$imd^{CA}$ flies were reared until 10-day-old. n=4 for all genotypes. Statistics: unpaired two-tailed Student's *t* test. Each graph shows the mean ± SEM.

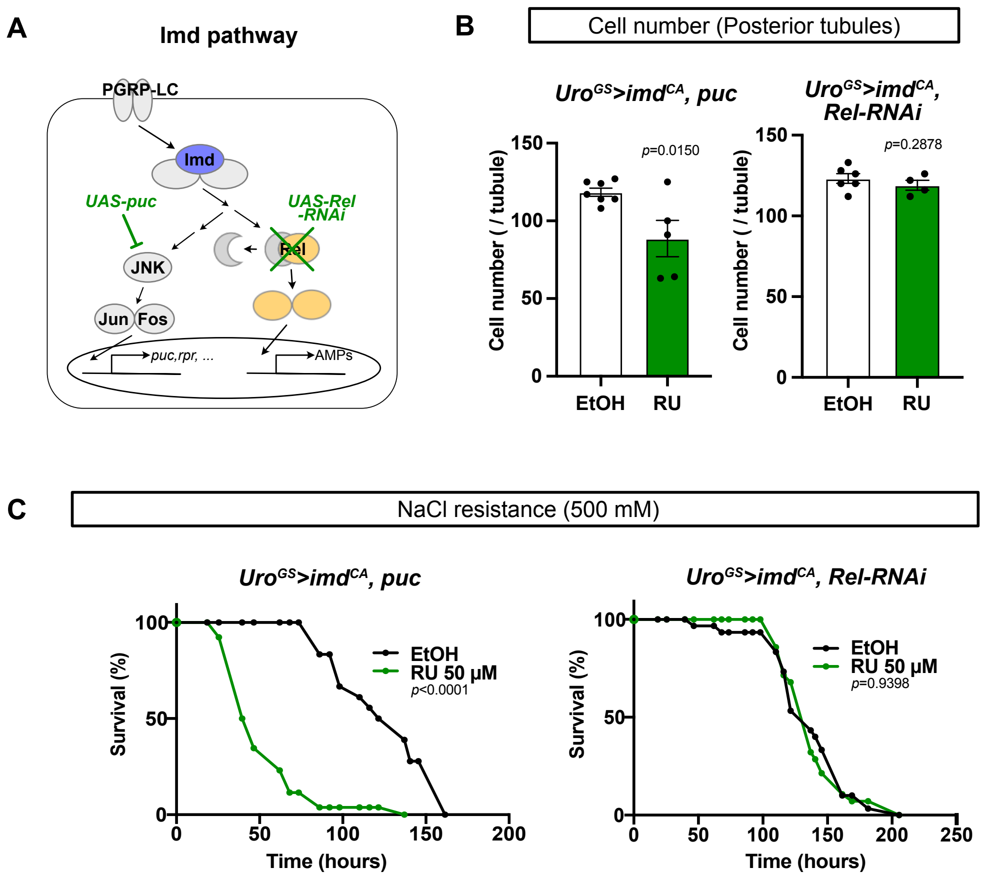


**Fig. S2 Tubular damage caused by Imd activation is *Relish* dependent.**

(A) A schematic image of the Imd pathway. (B) Quantification of the cell number of posterior tubules of female flies. n=7 for *Uro*$^{GS}$>*imd*$^{CA}$*, puc* EtOH, n=5 for *Uro*$^{GS}$>*imd*$^{CA}$*, puc* RU, n=6 for *Uro*$^{GS}$>*imd*$^{CA}$*, Rel-RNAi* EtOH, and n=4 for *Uro*$^{GS}$>*imd*$^{CA}$*, Rel-RNAi* RU. Statistics: unpaired two-tailed Student's *t* test. (C) 500 mM NaCl resistance of female flies. n=18 for *Uro*$^{GS}$>*imd*$^{CA}$*, puc* EtOH, n=26 for *Uro*$^{GS}$>*imd*$^{CA}$*, puc* RU, n=27 for *Uro*$^{GS}$>*imd*$^{CA}$*, Rel-RNAi* EtOH, and n=25 for *Uro*$^{GS}$>*imd*$^{CA}$*, Rel-RNAi* RU. Statistics: log-rank test. For (B) and (C), flies were fed a fly diet containing EtOH or 50 µM RU486 for 12 days before the experiments. Each graph shows the mean ± SEM.

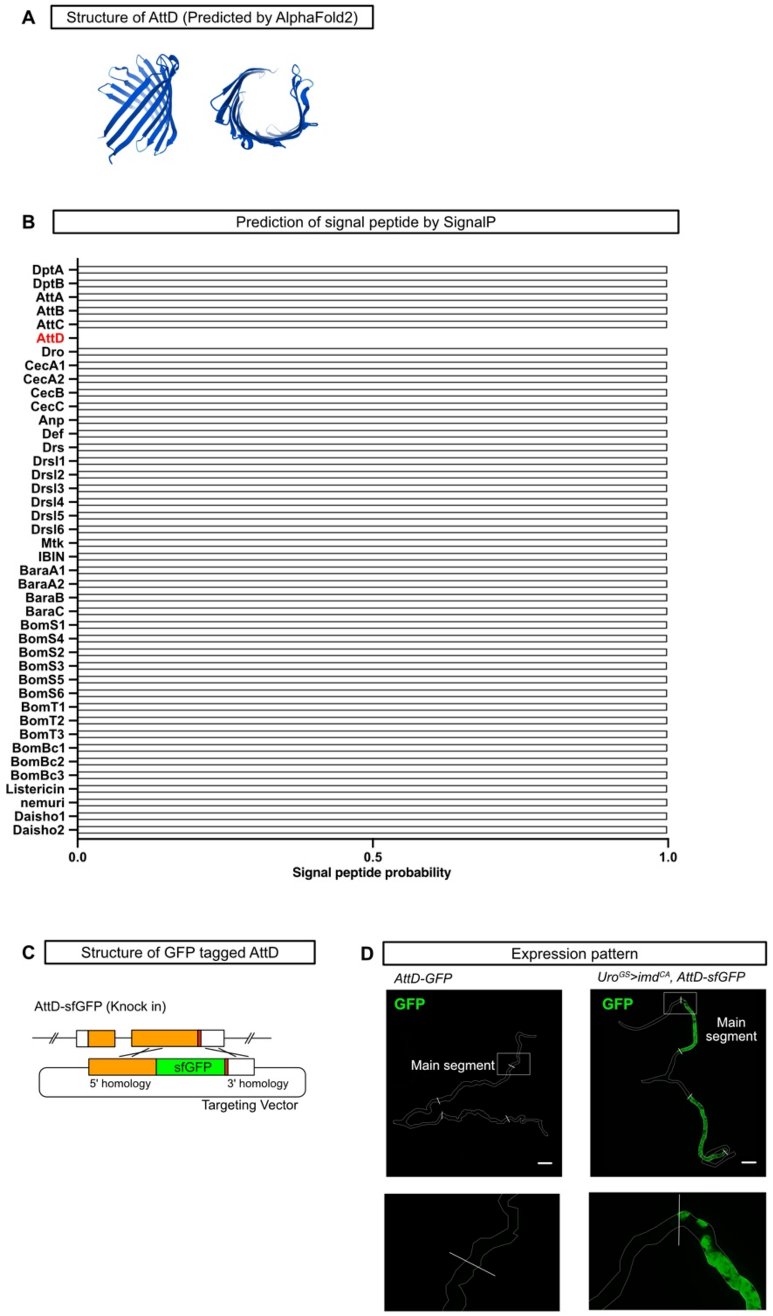


**Fig. S3 Attacin-D is one of the β barrel structure AMPs but cannot be secreted.** (A) The structure of AttD predicted by AlphaFold2. (B) Probability of a signal peptide of 42 AMP/AMP-like genes in *Drosophila melanogaster* calculated by SignalP6.0. See Table S3 for detailed results. (C) The structure of AttD-sfGFP (knock in) that we have generated using CRISPR/Cas9 system. Orange boxes show exons while bar shows an intron. Red box shows a stop codon. (D) Expression pattern of sfGFP knocked in AttD with or without Imd activation. UroGS >imdCA flies were fed a fly diet containing 50 μM RU486 for 3 days. Green: sfGFP signal (AttD). Scale bar: 200 μm. Lower panels are magnified images.

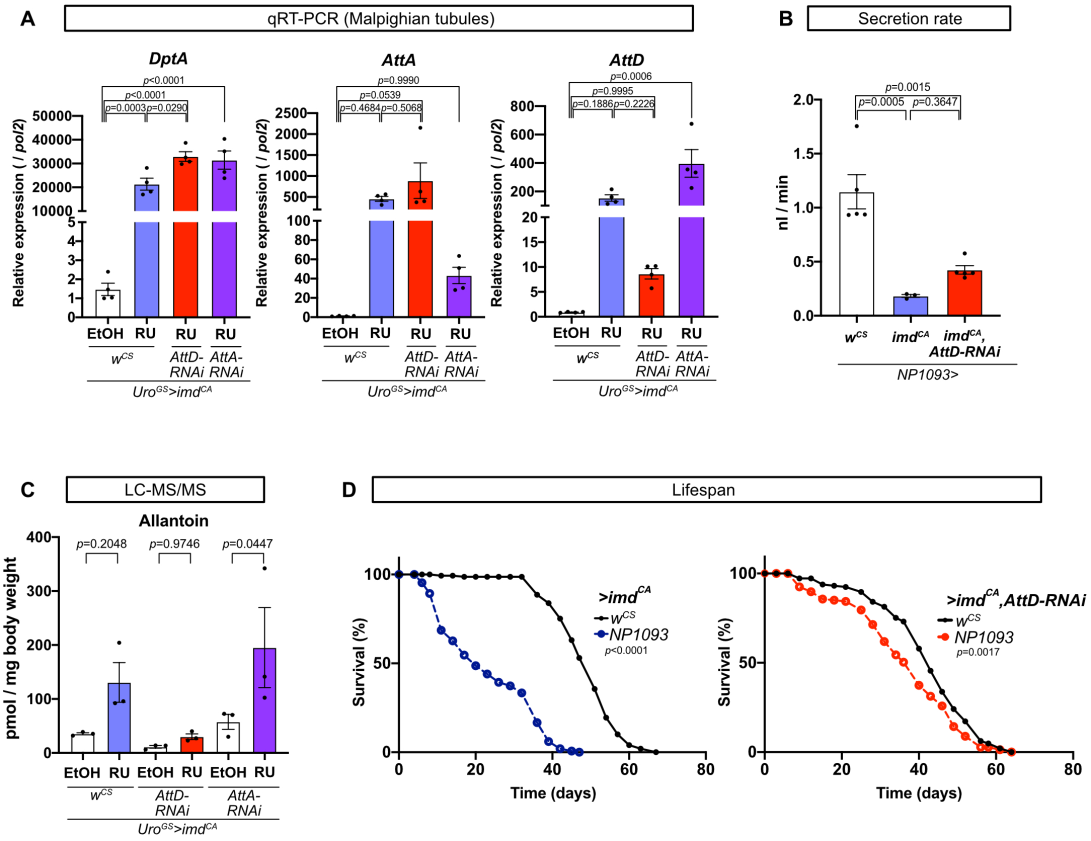


**Fig. S4 Knockdown of *Attacin-D* supresses Imd-induced renal damage without affecting other AMP expression.**

(A) Quantitative RT–PCR of the MTs of female flies. n=4 for all genotypes. Statistics: one-way ANOVA with Tukey's multiple comparisons. (B) Quantification of the tubular secretion rate of 10-day-old female flies measured by Ramsay assay. n=5 for *NP1093>w*$^{CS}$, n=3 for *NP1093>imd*$^{CA}$, and n=5 for *NP1093>imd*$^{CA}$*, AttD-RNAi*. Statistics: one-way ANOVA with Tukey's multiple comparisons. (C) Quantification of allantoin by LC-MS/MS in the whole body of female flies. n=3 for all genotypes. Statistics: one-way ANOVA with Sidak's multiple comparisons. (D) Lifespan of female flies fed a fly diet containing EtOH or 50 µM RU486. n=144 for *w*$^{CS}$*>imd*$^{CA}$, n=150 for *NP1093>imd*$^{CA}$, n=113 for *w*$^{CS}$*>imd*$^{CA}$*, AttD-RNAi*, n=125 for *NP1093>imd*$^{CA}$*, AttD-RNAi*. Statistics: log-rank test. For (A) and (C), flies were fed a fly diet containing EtOH or 50 µM RU486 for 12 days before the experiments. Each graph shows the mean ± SEM.

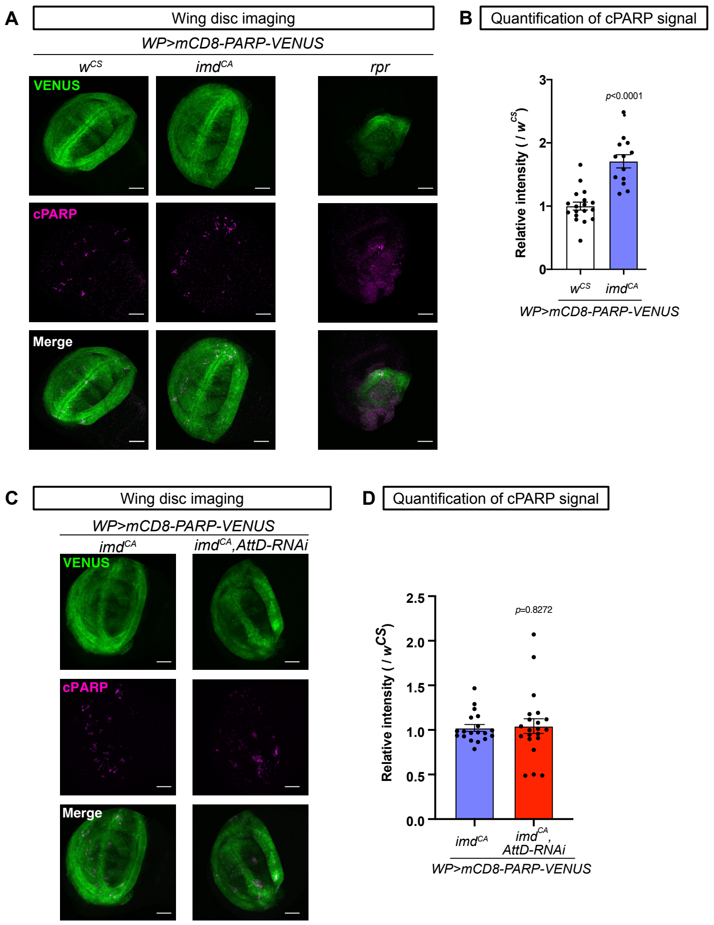


**Fig. S5 Attacin-D is not required for Imd-induced damage in wing discs.**

(A) Representative images of the stacked view of wing discs in larvae of each genotype. Green: VENUS signal, magenta: cleaved PARP (cPARP) signal Scale bar: 50 µm. (B) Quantification of the intensity of cPARP signal in wing discs. n=18 for *WP>mCD8-PARP-VENUS, $w^{CS}$*, n=13 for *WP>mCD8-PARP-VENUS, $imd^{CA}$*, and n=16 for *WP>mCD8-PARP-VENUS, AttD*. Statistics: one-way ANOVA with Tukey's multiple comparisons. (C) Representative images of wing discs in larvae of each genotype. Green: VENUS signal, magenta: cleaved-PARP signal. Scale bar: 50 µm. (D) Quantification of the intensity of cPARP signal in wing discs. n=13 for *WP>mCD8-PARP-VENUS, $imd^{CA}$*, and n=16 for *WP>mCD8-PARP-VENUS, $imd^{CA}$,AttD-RNAi*. Statistics: unpaired two-tailed Student's *t* test. Each graph shows the mean ± SEM.
